# Subjective Beliefs In, Out, and About Control: A Quantitative Analysis

**DOI:** 10.1101/2020.05.27.115998

**Authors:** Federico Mancinelli, Jonathan Roiser, Peter Dayan

## Abstract

A critical facet of adjusting one’s behaviour after succeeding or failing at a task is assigning responsibility for the ultimate outcome. Humans have trait- and state-like tendencies to implicate aspects of their own behaviour (called ‘internal’ ascriptions) or facets of the particular task or Lady Luck (‘chance’). However, how these tendencies interact with actual performance is unclear. We designed a novel task in which subjects had to learn the likelihood of achieving their goals, and the extent to which this depended on their efforts. High internality (Levenson I-score) was associated with decision making patterns that are less vulnerable to failure, and at the same time less oriented to more rewarding achievements. Our computational analyses suggested that this depended heavily on the adjustment in the perceived achievability of riskier goals following failure. We found beliefs about chance not to be explanatory of choice behaviour in our task. Beliefs about powerful others were strong predictors of behaviour, but only when subjects lacked substantial influence over the outcome. Our results provide an evidentiary basis for heuristics and learning differences that underlie the formation and maintenance of control expectations by the self.

## 1. Introduction

The controllability of a particular environment quantifies the extent to which the actions that we can exert in that environment take us adequately quickly, reliably and cheaply to desired outcomes, such as the acquisition of rewards and the avoidance of punishments. Uncontrollable environments may be temporarily benign – but for our reactions to new opportunities and threats to be successful, controllability is essential.

Versions of this objective notion of controllability underpin work in areas such as learned helplessness (when animals learn that one environment is uncontrollable, and generalize this to new situations; Seligman & Maier, 1967). Learned helplessness is pervasively used as a model of depression (see, e.g. Sherman et al., 1982), and attempts have also been made in the field of reinforcement learning to look in a more granular manner at formalizing the separate components of controllability in terms of prior expectations (Huys & Dayan, 2009).

However, it is the subjective perception of controllability that has the ultimate psychiatric import. This perception can diverge substantially from the objective facts, particularly in the way it affects the ascription of responsibility for successes and failures (Miller & Seligman, 1975; Klein et al., 1976; Kuiper, 1978). People who ascribe failure to the vicissitude of bad luck or evil underlying forces, might predict better future outcomes than people who think that their internal incompetence is at fault, and believe they will require minor degrees of behavioural adaptation to obtain better performances. Moreover, from an emotional standpoint, they might experience more frustration and stress having to face a punishment they felt was to some degree unavoidable (Julian et al., 1968). This subjective perception of controllability, which in fact individuates a subjective causal model for the experience of events, is captured by scales such as the locus of control (LoC; Rotter, 1966). Such ascriptions have an important, and complex knock-on effect on learning, since, for instance, if chance can be blamed for failures, and skill for successes, subjects will build a severely biased impression for themselves of ‘objective’ controllability.

Most human studies attempting to individuate the behavioural implications of LoC and attribution have done so by having different subject groups fill in questionnaires and measuring difference in their responses (e.g. Levenson, 1973), or framing tasks as being determined by skill/chance (Rotter, 1966; Phares, 1957; James & Rotter, 1958; Rotter et al., 1961), or using simple betting, or risk-aversion paradigms (e.g. Liverant & Scodel, 1960). Thus, we lack a task that decomposes objective controllability into its finer parts described above, and assesses the interaction between outcomes, ascription style, expectations and choices.

In our task, subjects choose between and use one of a range of tools to achieve various simple goals. The tools are only partially reliable (with differing amounts of entropy in the effects of choices) and intermittently effective (affording many or few actions per unit interval; a quantity we call influence), to degrees that need to be learned. In turn, subjects learn about the resulting achievability of rewards. This allows us to examine the interactions among prior expectations, attributions, and learning, on performance.

Our primary finding is that, in both conditions of high and low contingency between effort and outcome, internality with respect to self (Levenson I-scores; Levenson, 1973), was inversely correlated with choices of less attainable, more rewarding goals. In good influence conditions, where efforts are predictive of outcomes, internal subjects prioritized features that are tied to achievability, and in doing so managed to lose on fewer trials. In both good and poor influence conditions, internality was strongly tied to the speed of learning from failure, that the more rewarding goals would not be achievable. High I-scorers learned unachievability faster, whilst low I-scorers perseverated in attempting the next-to-impossible, more rewarding goals. Lastly, we found serendipitously that beliefs about powerful others were strongly predictive of performance in low influence conditions.

## 2. Methods

### 2.1. Participants

We recruited 37 subjects through the Institute of Cognitive Neuroscience subject database at University College London (21 females; age, median : 24, highest : 39, lowest : 18 yrs). All subjects completed a structured telephone interview to confirm the absence of any previous or current mental disorder. On arriving in the lab, subjects approved and signed an ethics consent form, and were instructed to read a briefing sheet which explained the task. Before starting the task fully, subjects performed 3 practice trials in both high and low influence conditions (see Experimental design), under supervision of the experimenter; they were told that they could ask for clarification if they were unsure about any aspect of the experiment, and that they could begin when they were ready to do so. After performing the task, subjects completed Levenson’s LoC questionnaire (Levenson, 1973). They then collected a payoff which consisted of the sum of £10 and the actual outcomes of 30 trials sampled at random. Subjects knew the structure of the payoffs. Two female subjects were excluded when performing data analyses. One subject was excluded on account of having too many irrational vehicle and goal choices on *Catch* trials (see the table in figure 1) (more than 4 standard deviations away from the mean of the remaining subjects). This may be indicative of poor understanding or lack of focus throughout the task. A second subject was excluded due to an outlying low I-score (more than 3 standard deviations away from the mean of the remaining subjects). These exclusions left us with 35 subjects.

**Figure 1:**
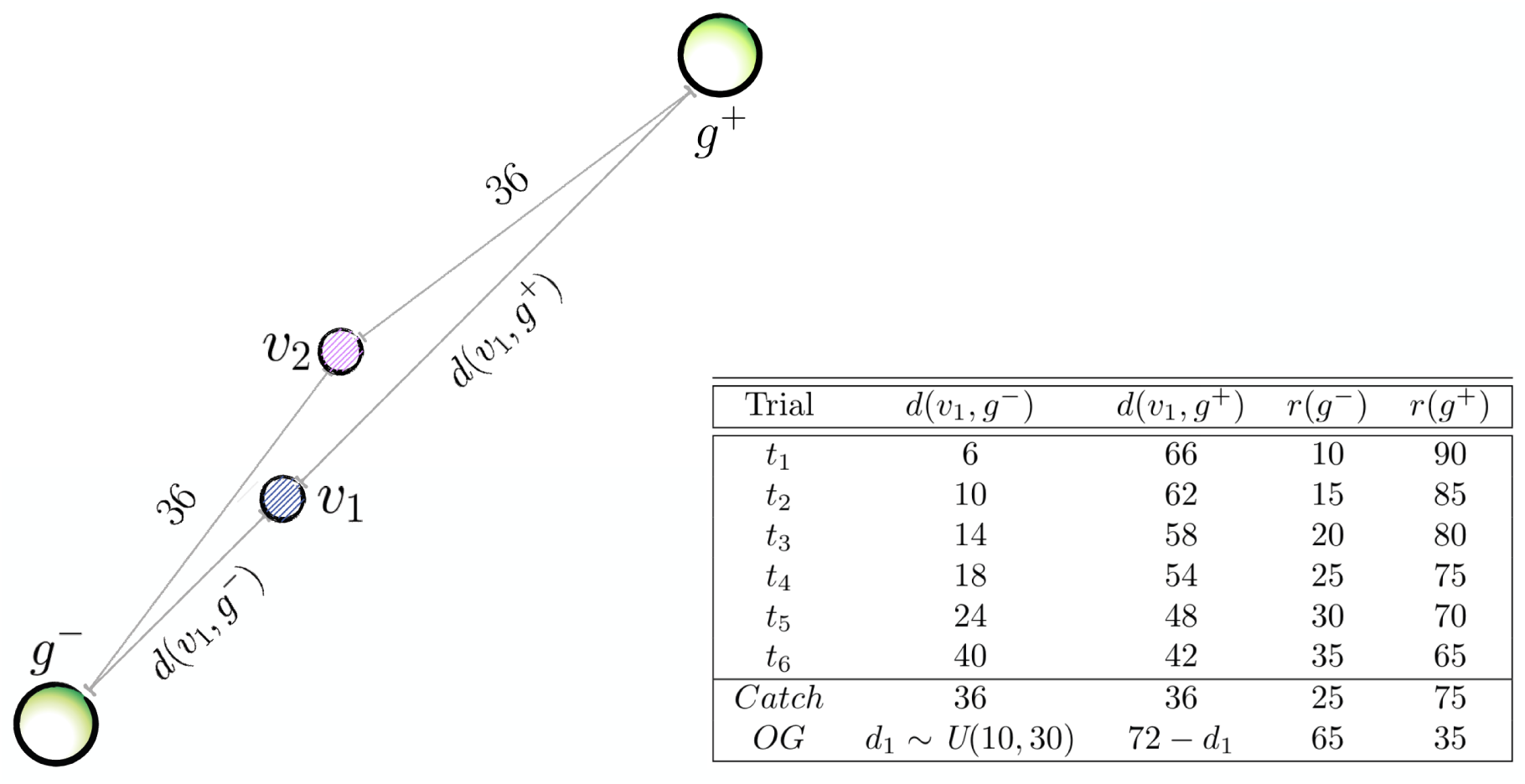
Illustration of double-goal trials (DGTs). Vehicles’ colours in this figure are random. *d*(*v*_1_, *g*^−^) and *d*(*v*_1_, *g*^+^) indicate the Manhattan distances (in units of the vehicles step size) from vehicle *v*_1_ to both *g*^−^ and *g*^+^. Vehicle *v*_2_’s distance from both goals is constant, i.e. *d*(*v*_2_, *g*^−^) = *d*(*v*_2_, *g*^+^) = 36. The last two columns indicate the monetary reward (in pence) associated with each goal, i.e. *r*(*g*^−^) and *r*(*g*^+^). The theme underpinning the design is, the closer a vehicle is to *g*^−^, the less rewarding *r*(*g*^−^) will be, and the more rewarding *r*(*g*^+^) will be. In this illustration, we arbitrarily placed *v*_1_ asymmetrically (closer to *g*^−^), and *v*_2_ at the center of the canvas. Recall, however, that all trials showed in the table appear twice in every block, the second time with vehicles positions swapped. This was done to ensure that we can fully orthogonalise rewards, distances and guidabilities. Thus, if we first encounter trial *t*_4_ with *v*_2_ at the center of the canvas (as is shown), the second time we meet this trial type the vehicles’ positions will be swapped (and the canvas randomly rotated) and *v*_2_ will be the one closer to *g*^−^. *Catch* and *OG* trials are atypical, and test participants’ notion of the vehicles’ guidabilities, and break the inverse correlation of distance and reward imposed by the design of the other trials, respectively. In *Catch* trials, vehicles are equidistant from both goals, so we expect participants to choose the more guidable vehicle (*v*_1_) and the more rewarding goal. Obvious-goal (*OG*) trials are the only occasion in which *r*(*g*^−^) is actually larger than *r*(*g*^+^), so that choice should “obviously” be *g*^−^. The distance of the closer vehicle to the 65p goal is extracted from a uniform distribution in the indicated set (we used a MatLab notation to denote the sample space); this is the only occasion in which distances are sampled randomly within the design. All DGTs are individually rendered, for clarity, in Supplementary Materials, where we show a full instance of the first 16 trials in a block. An *OG* trial is also shown in figure 4.

### 2.2. Questionnaire

The perception of behavioural control is traditionally measured using questionnaires. Rotter (1966) introduced the I-E (Internal-External) forced-choice scale. Here, we adopted a more finely graded, Likert type scale (Levenson, 1973). This dissects LoC into three separate dimensions: Internality (I), Chance (C), and Powerful Others (P). In her work, Levenson motivates this choice: “The rationale behind this tripartite differentiation stemmed from the reasoning that people who believe the world is unordered (chance) would behave and think differently from people who believe the world is ordered but that powerful others are in control. […] Furthermore, it was expected that a person who believes that chance is in control (C orientation) is cognitively and behaviourally different from one who feels that he himself is not in control (low I scale scorer)”.

In a further difference from Rotter’s original scale, the statements underpinning Levenson’s assessments are phrased so that it is clear that they pertain only to the person answering. That is, they measure the degree to which an individual feels she has control over what happens, rather than what she feels is the case for people in general. Levenson (1981) also ensured that the LoC measures were not influenced by demand characteristics, showing that there was no significant correlation between her scores and the Marlowe-Crowne Social Desirability Scale (Crowne & Marlowe, 1960).

Table 1 shows empirical cross-correlations between the different items found in our dataset. These results are consistent with the literature (Levenson 1973).

**Table 1:** Intrinsic LoC scores correlations. None reaches significance at 0.05.

|  | Internality (I) | Chance (C) | Powerful others (P) |
| --- | --- | --- | --- |
| I | - | $r = -.24, p = .16$ | $r = -.29, p = .08$ |
| C | - | - | $r = +.29, p = .08$ |
| P | - | - | - |

### 2.3. Power analysis

Our analyses aim at resolving the individual relevance of LoC subscales in our task, in which we hypothesise Levenson I- and C-scales to play substantial roles. With 35 subjects, we had a power of 0.8 to detect correlation coefficients of ∼ 0.45, with a two-tailed significance threshold of *p* = 0.05.

### 2.4. Experimental design

The task consists of a novel videogame (coded in JavaScript). On each trial, subjects see a large blank canvas, and have to select between, and then move, one of two vehicles (shown as small circles) to a goal (large circle) within 14 seconds by pressing the arrow keys on a keyboard. Movement is only allowed horizontally and vertically, and in predefined step-sizes. If subjects are successful, they win a reward whose magnitude (in pence) is signalled at the goal. If they fail, they lose a fixed amount of £0.15. Vehicles differ in their placement on the canvas, and in the extent of control they afford (guidability). A vehicle’s guidability simply defines how likely it is to move in the direction implied by the arrow key pressed by the subject as opposed to moving in any cardinal direction (chosen apparently uniformly at random). Subjects learn about the characteristics of each vehicle from the experience gathered within a block. On some trials, subjects are offered two potential goals; on others, there is only one. Goals differ by the amount of reward, and, in most trials with more than one goal, the one that is more rewarding is further away than the one that is less rewarding. Trials differ in registering more (high influence) or fewer (low influence) actual moves per second, though subjects can always press keys as fast as they like. In order to avoid potentially confounding fatigue effects, participants are offered the opportunity to rest at the end of every trial (by adding 10 seconds to the inter-trial interval) before the next trial starts. Figure 4 illustrates the timeline of a trial with two goals.

**Figure 2:**
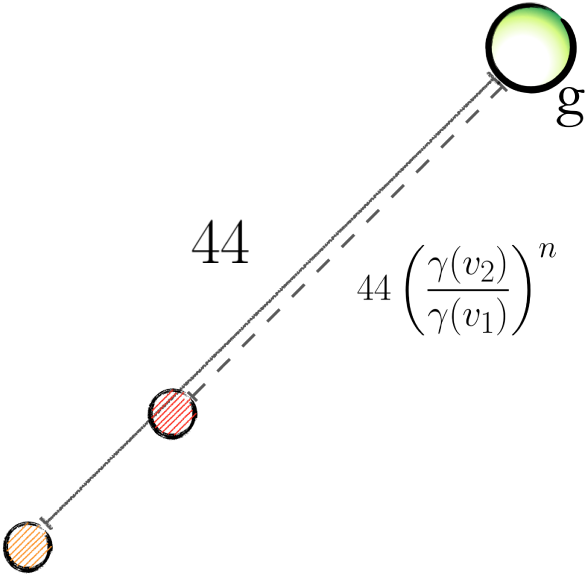
Illustration of single-goal trials (SGTs). Vehicle’s colors in this figures are random. In SGTs, the vehicles’ relative positions from the goal (*g*) depends on their guidability. The more controllable (yellow, in figure) vehicle’s distance from *g* is always 44 Manhattan steps. The other (red) vehicle’s distance is instead a fraction of this distance equal to powers (the exponent *n* takes the values 1,2,3 and 4) of the ratio of the guidabilities, i.e. *γ*(*v*_2_)*/γ*(*v*_1_). Setting the distance of the red vehicle from *g* to be dependent on this ratio promotes harder decision making; and we empirically verified that it ensures that *v*_1_ is the correct choice exactly half of the time if subject pressed at a frequency of 8*Hz* (in high inuence conditions, it is best to choose the most controllable vehicle when *n* < 3, and the closer vehicle when *n* > 2). Thus, choosing the same vehicle on all four SGTs is then wrong in conditions of high inuence. If the two vehicles are equally controllable, their distance from the goal is randomly 44 and 38 Manhattan steps, and of course the closest vehicle to *g* is always the right choice. The probability of making the goal in low influence conditions is instead hopeless, by design (no win was recorded). Rewards associated with the single goal are pseudorandomly distributed in a uniform interval between 50 and 80 pence, in steps of 10. The order of the four SGTs is randomized,, but they always occur at the end of each block, when the guidability of each vehicle can be assumed to be known by participants.

**Figure 3:**
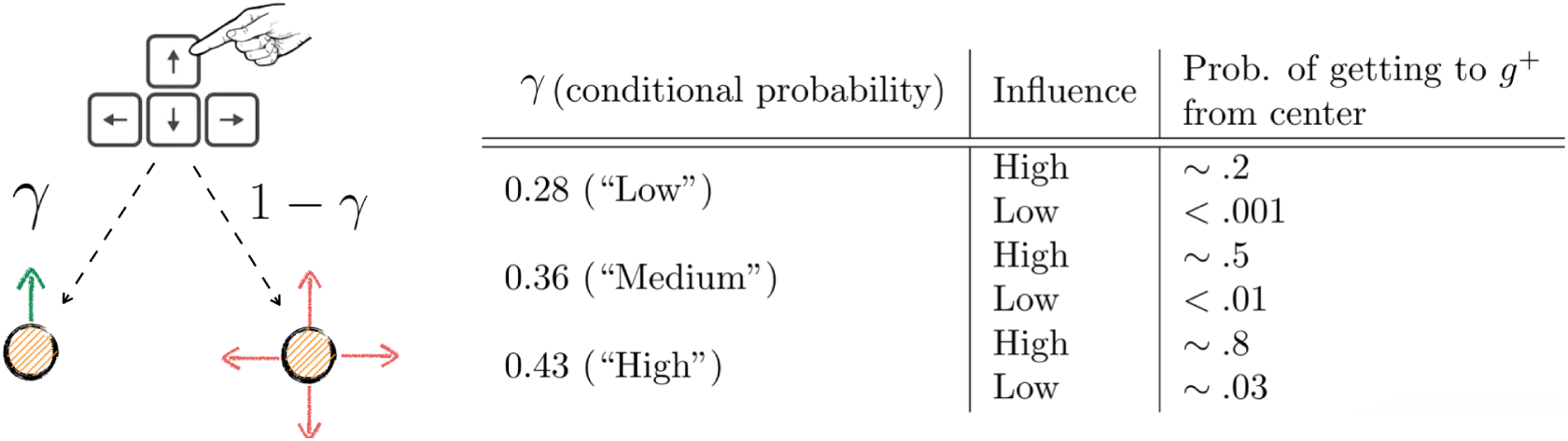
The design of guidability. The figure on the left hand side depicts the relationship between miroscopic actions and outcomes with respect to the conditional probability *γ*. On pressing (for instance) the upwards arrow key, the vehicle (small yellow circle) follows the direction verbatim with probability *γ* (left; green arrow) or goes in a direction sampled uniformly at random (right; red arrows). On the right hand side, a table describes what the three different values for *γ* imply when a vehicle placed at the center of the canvas is set to reach *g*^+^, according to whether influence conditions are high or low. Reaching *g*^+^ from the center of the canvas in low influence conditions is almost impossible.The choice of guidabilities implied substantially different probabilities of reaching a goal from the central position (using the central vehicle) in high influence blocks. For instance, pressing consistently at a frequency of 8*Hz* for the full duration of the trial (14s) would yield probabilities 0.8 (“High”), 0.5 (“Medium”), and 0.2(“Low”). In low influence blocks, the chance of making a goal using the central vehicle was circa 0.03 (“High” guidability) and fell substantially below 0.01 when guidability is “Medium” and “Low”. These numbers are Monte Carlo estimates obtained via empirical simulations, which assumed an optimal strategy of pressing, consisting of moving the vehicle only in the direction of the goal in the allotted time, at the maximum frequency allowed (8 or 4 Hz according to influence condition). Note that while we used empirical probability of reaching the goal from the central position to inform the value of guidability, we did not use it to determine the relative positions of the other vehicle.

**Figure 4:**
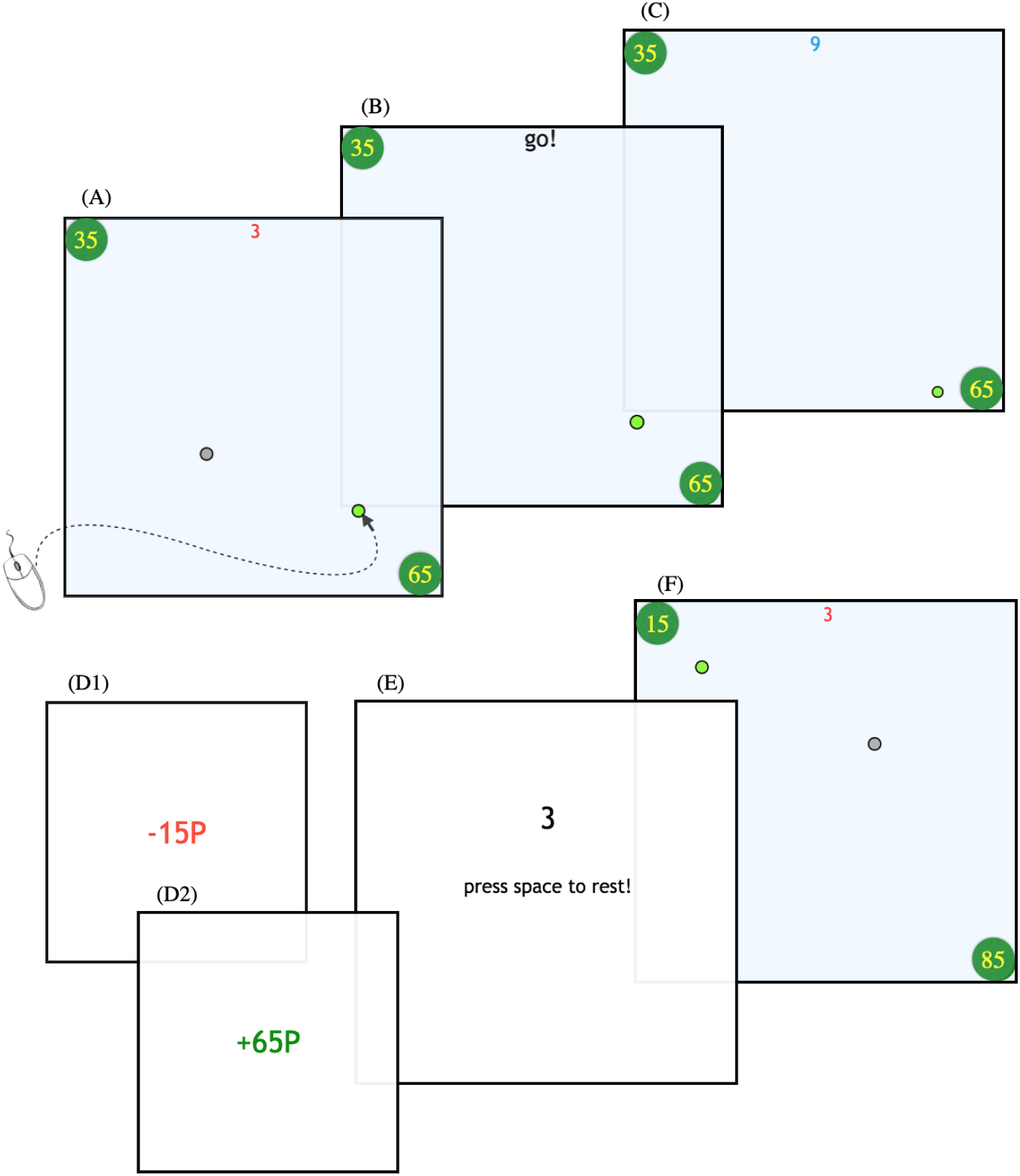
The timeline of a trial. This trial is of type *OG* (see figure 1). This is the only trial type where *r*(*g*^−^) is larger than *r*(*g*^+^), intended to prevent subjects from starting to assume that the larger goal would always be further away in DGTs. From top-left to bottom-right: (A) vehicle selection phase. Subjects had 5 seconds (3 seconds of which are left in the figure) to choose a vehicle using the mouse (whose movement is shown by the curved arrow). The ultimate goals are shown as green circles whose values in pence are shown as numbers inside each. (B) Any unchosen vehicle is removed, and a go! cue starts the movement phase of the trial. The subject then had 14 seconds to drive the chosen vehicle to reach one of the goals to win the amount shown. Otherwise, they lost 15p. (C) the trial is in progress - in this case with 9s left (top centre) for moving. (D1) shows what subjects see upon expiry of the allowed time; the amount lost (15p) is displayed in red, and is accompanied by a noxious sound. (D2) shows what subjects see if they touched the goal before the time elapsed. The amount won is displayed in green accompanied by a pleasurable sound. (E) waiting for the next trial to start (3s). At the end of the twentieth trial of each block, participants saw a sign saying “vehicles change”, which notified the end of the block. They then had a 5s rest before starting again. On pressing the space bar, 10s would be added to the count-down. (F) The new trial (of type *t*_1_) begins.

#### 2.4.1. Control manipulations

We manipulated the probabilities of reaching the goal in three ways. The first, and most significant, is the *distance* from each vehicle to each goal. The second was the *guidability* of each vehicle, which required learning. We parameterized the guidability of the vehicles in terms of conditional probability (Huys & Dayan, 2009): independently, at each press, the vehicle follows the subject’s intended direction (i.e., the chosen arrow key) with probability *γ*, or moves in a random direction (drawn uniformly) with probability 1 − *γ*. Note that this includes the arrow direction followed by chance. Finally, perhaps the most restrictive manipulation of achievability was a hard limit on the maximum number of steps per second that vehicles would move. We signaled two types of block: a benevolent type (“high influence”) in which this limit was set to 8 steps per second (8*Hz*), and a malevolent type (“low influence”) in which this limit was set to 4*Hz*. Note that all subjects could reach (and in many cases, break) the maximum pressing frequency of 8*Hz*. In high influence blocks, higher pressing frequencies are designed to entail higher rewards, but in low influence blocks, the two are uncorrelated. We introduced this manipulation to test whether influences of internality on behaviour would be more salient in inherently good or bad environments. The influence manipulation is conceptually different from that of vehicular guidability: it limits the total influence that the subject has on the microscopic outcome (moving vs. not moving), rather than the noise in the microscopic action-to-outcome link (moving in the intended vs. random direction). In order to maximize rewards, subjects should take all three factors into account: large distances could be covered quickly with a good vehicle, just as small distances could be hard to cover, if the vehicle chosen is not very guidable and/or the block is low influence. Collectively, the three manipulations of distance, guidability, and influence provided for an extensive range of levels of achievability.

#### 2.4.2. Task structure

The full task comprised 160 trials, divided in 8 blocks. Within each block, the first 16 trials included two goals (double goal trials; DGTs). Four single goal trials (SGTs) appeared at the end of each block, by which time we assumed that subjects would have a good grasp of the controllability of each of the two vehicles. The design for both DGTs and SGTs is illustrated in figures 1 and 2. The influence condition, or block type (“high” or “low” influence) was manifest in the background colour of the canvas, either light beige or blue (counterbalanced across subjects). Subjects were instructed as to the link between background color and influence.

DGTs (128 trials per subject) always included two vehicles and two goals. Here, subjects could decide between plans offering different tradeoffs of achievability, both in the form of distance and vehicle guidability, and reward. DGTs allow us to examine learning of achievability and its components throughout a block. At the start of each block, subjects have no knowledge about the worth of each vehicle, and must learn in order to know what they can, and cannot, do. All blocks involved all the DGTs (8 trial types) indicated in the table to the right of figure 1 in random order. The same trial appears twice, (1) with a randomly rotated canvas (by multiples of 90 degrees), and (2) with vehicles swapping positions; this is in order to obtain a fully orthogonal design. In DGTs, there is always a vehicle at the center of the canvas, so that subjects face the recurrent question, at each trial, of whether they will be able to achieve the more rewarding goal using the vehicle currently at the center; as we will see, our computational analyses will shed some light on the learning process underlying the formation of this judgement. In Supplementary Materials, we illustrate an instance of how the 16 DGTs might be laid out in a block.

SGTs (32 trials in total per subject) were explicitly designed to pit goal distance against vehicle guidability, to reveal whether subjects might exhibit a bias towards either, or whether they would choose approximately optimally, according to the probability of reaching the goal. SGTs were, on average, harder in both influence conditions. This is because the furthest vehicle from the single goal is at a distance larger than that between the central vehicle and the more rewarding goal in DGTs. Throughout SGTs, the more guidable vehicle is always at a distance of 44 Manhattan steps from the goal; the less guidable one lies instead at a closer distance to the goal which is a direct function of the fraction of the vehicles’ guidabilities. This heuristic was used to guarantee that in two out of 4 SGTs, the best vehicle to use would be the closest to the goal, and in the reamaining two, the farthest.

Within each influence condition, the four blocks differed in the guidability of the pair of available vehicles. In one block, the two vehicles would be equally guidable, and their guidability level would be “Medium” (see figure 3 for the values of guidability of each level); in the remaining three, they would be faced with a (“Medium”, “High”) pair, a (“Low”,”Medium”) pair, and finally a (“Low”,”High”) pair. Vehicles could always be identified through their colours within a block, but their associated guidabilities had to be learnt from the start of each new block. The design choices of the conditional probabilities to assign to vehicles were based on the fine tuning of the empirical odds of reaching *g*^+^ from the central position of the canvas (empirical simulations described in Supplementary Materials). This process produced values for *γ* which were used during the main experiment as per figure 3.

#### 2.4.3. Notation

Throughout the text, we denote the two vehicles in a block as simply *v*_1_ and *v*_2_. Recall that vehicles are identified by their colours, and differ in their guidability values (except for one block per influence condition where both of them have “Medium” guidabilities). We denote vehicle *v*_*i*_’s guidability (as inferred by trial *t*) as *γ*_*t*_(*v*_*i*_), and its true guidability as *γ*(*v*_*i*_). As SGTs are faced after 16 trials in each block, we drop the pedix *t*, and safely assume subjects know the true guidability of the vehicles. Without loss of generality, we will always assume that *v*_1_ is the more guidable vehicle. We will specify that *v*_1_ and *v*_2_ are equally guidable when this is the case (i.e. both have “Medium” levels of the guidability). Finally, again in DGTs, we will use an auxiliary variable (*c*_*t*_) to indicate which vehicle is at the center of the canvas at trial *t*, so that, for instance, *c*_*t*_ = 1 indicates that *v*_1_ is at the center of the canvas. Recall that each of *v*_1_ and *v*_2_ is placed at the center of the canvas an equal number of times through the DGTs in a block.

A goal will simply be denoted *g*, and *r*_*t*_(*g*) the reward connected to it at trial *t*. We will add an apex as a shortcut to signalling that one goal is the more rewarding one in a DGT, so that *g*^−^ signals the less, and *g*^+^ the more rewarding goal. For SGTs, where there is only one available goal, we will simply write *g*.

### 2.5. Model agnostic analyses

We examined the relationship between I- and C-scores and our outcome measures. We had no hypotheses concerning P-scores, so exploratory analyses on these data are reported separately. All our model-agnostic analyses were performed in each influence condition separately.

In DGTs, maximising rewards requires learning the achievability of goals via using the vehicles available in the block at hand. Nevertheless, choosing the more rewarding goal (*g*^+^), regardless of the block we are in or which vehicle is selected, is always by design less likely to lead to success (percent success of reaching *g*^+^, from data: 45% in high influence blocks, 3% in low influence blocks; of reaching *g*^−^: 67% in high influence blocks, 43% in low influence blocks). Thus, the proportion of attempts on *g*^+^ goals is a direct readout of the subjective trade-off between achievability and reward, and will be a key outcome measure.

We report the proportion of *g*^+^ attempts made using the more guidable vehicle (*v*_1_), which we chose as a measure of sensitivity to control, functional to reward. This measure of course depends on the guidability inherent to *v*_1_ (i.e. *γ*(*v*_1_)). Finally, we report basic performance metrics such as the amount of money accrued and the number of losses, and examine how these are affected by control expectations as measured by LoC. Two trials in every block are catch trials (type *Catch* in the table in figure 1); these see both vehicles placed at the center of the canvas and make it obvious that the choice should be of the most controllable vehicle (and pursuing *g*^+^). We are interested in the proportion of correct choices here as a proxy for how important vehicular controllability was for a subject. In single goal trials, we just measure the trade-off between sensitivity to guidability and distance.

### 2.6. Model dependent analyses

Our model-dependent analyses were intended to provide a clearer view over the model-agnostic effects observed in raw data. We employed a hierarchical graphical structure for all our models (Daw et al., 2011). All models contained various parametrized additive factors intended to estimate a net value of choosing a combination of vehicle and goal, given the features of the trial at hand. Posterior distributions over the parameters for each model for each participant and condition were estimated using a Hamiltonian Monte Carlo sampler, implemented in Stan (http://mc-stan.org; Carpenter et al., 2017). Model assessment was performed via leave-one-subject-out cross validation (LOSO), obtaining average log-likelihoods for each subject as they were held out of training and only used as test data. Models were compared by assessing the distribution of the inter-model difference in log-likelihoods for each trial for each subject and considering whether this differed significantly from zero and with what sign. We also generated synthetic data from our winning models to assess the fidelity with which they could recapitulate the statistical characteristics of the original behavior (see Supplementary Materials).

Statistical analyses related to LoC were based on permutation tests with empirical null distributions created by randomizing questionnaire scores across subjects. In addition to considering the relationship between parameters obtained and questionnaire scores, we tested whether the inferred parameter fits were *necessary* to recover the main model-agnostic effects observed in the raw data. For each parameter, this was done by (1) permuting its estimates across subjects, (2) generating synthetic data under the permutation (1000 data-sets), and (3) considering how many of the synthetic effect sizes, in the null distribution thus obtained, exhibited a larger magnitude than in the original labelling.

Finally, in order to establish the importance of recovered parameter estimates for prediction of individual LoC scores, we harnessed measures of conditional variable importance in the context of regression through random forests (Strobl et al., 2008). The rationale is the same as in the previous approach: a variable is shuffled and utilised in regression in a random forest, and the resulting mean squared error is compared with the value found with the original labelling.

The features of the trial that determined the additive components of the estimated value included vehicle-goal distance, vehicular guidability and reward size. These make up the value of choosing a vehicle-goal pair, which then generates decisions according to a softmax policy. Influence conditions were considered separately, with each including its own set of parameters. We added extra components to model behavioural change across the whole task.

For double goal trials, we incrementally built a sequence of increasingly more sophisticated models. We started from (1; “Additive”) additive influences of vehicle-goal distance, vehicular guidability and reward size; and leading through (2; “Bias”) a bias term indicating an absolute propensity to choose (or avoid) *g*^+^, irrespective of the characteristics of the trial or any temporal aspect; (3;”Temporally evolving bias”) a temporally evolving bias which embodies a propensity to choose *g*^+^ which diminishes (perhaps as a consequence of fatigue) or increases, in time; (4; “Win-stay Lose-shift”) a term which modelled the propensity to choose either vehicle to attain *g*^+^, which varied as a function of the outcome of the previous trial, (5; “Vehicle independent RW”) a Rescorla-Wagner term which promotes or discourages the choice of *g*^+^ with either vehicle, by keeping track of a running average value of choosing *g*^+^ (this conflates both vehicles into one value); and finally (6;”Vehicle dependent RW”) a different Rescorla-Wagner term which does the same, but keeps separate records for the two vehicles. Note that, in DGTs, we do not consider interactions between predictors, as these would be too many to individually test.

For single goal trials, we assumed that subjects employed a policy based on (1) additive influences of vehicular guidability and vehicle-goal distance; (2) as in (1), but adding an interaction term of guidability and distance; (3) a combination of guidability, distance, and the pure probability of reaching the goal with either vehicle (the computation of which is shown in Supplementary Methods); or, finally, (4) the pure probability. It is important to note that all these formulations amount to different approximations of the computation of probability; arguably, from the simplest, i.e. (1), to the most complete, (4).

Formal descriptions of the models along with their equations can be found in Supplementary Materials.

## 3. Results

### 3.1. Model agnostic results

#### 3.1.1. Validity of control manipulations

We first assessed whether our control manipulations worked as planned. As intended, individual trial-averaged pressing frequencies were a strong predictor of performance in high influence blocks (money earned: *r* = .71, *p <* 0.001, 95%CIs [.50, .85]; num. of losses: *r* = −.64, *p <* 0.001, 95%CIs [-.80, -.40]). Conversely, due to the cap at 4*Hz*, trial-averaged individual pressing frequencies were not significantly predictive of performance in low influence conditions (money earned: *r* = .20, *p* = .24, 95%CIs [-.14, .50]; num. of losses: *r* = −.13, *p* = .44, 95%CIs [-.44, .20]). Thus, by design, half of the blocks arranged a contingency between effort (in the form of pressing frequency) and reward, while the other half did not. Finally, as we hoped, vehicles’ controllabilities could be discriminated quite reliably by the 16^th^, *Catch* trial (where both vehicles are placed at the center of the canvas), with a choice frequency of the more controllable vehicle averaging at 71% (high influence) and 73% (low influence), a good indication that by the time subjects were faced with SGTs, they generally knew which vehicle was most controllable.

#### 3.1.2. Double goal trials

We found that the proportion of *g*^+^ attempts was anticorrelated with I-scores in both high (*r* =. − 50, *p* = .002, 95%CIs [-.71, -.20]) and low influence conditions (*r* = −.55, *p <* .001, 95%CIs [-.75, -.27]) (see figure 5); contrary to our predictions, we found no such relationship with respect to C-scores (high influence: *r* = − .09, *p* = .61, 95%CIs [-.41, .25]; low influence: *r* = .20, *p* = .26, 95%CIs [-.14, .49])). The vast majority of attempts to *g*^+^ goals were made using the central vehicle (92%, in both influence conditions).

**Figure 5:**
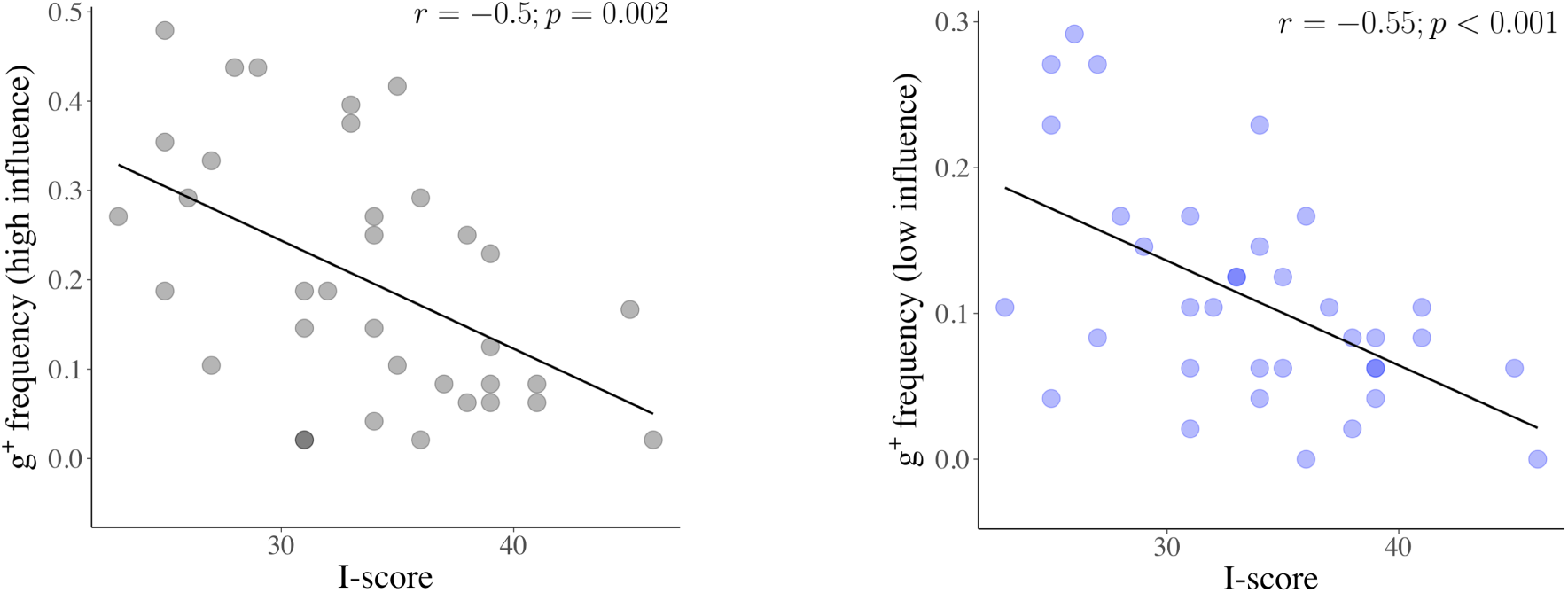
The negative correlations between I-scores and choice frequency of *g*^+^ (riskier, more rewarding) goals, in high influence (gray dots; left) and low influence (blue dots; right) conditions. Note the differing range of values (on the y-axes) which reflect the overall reduced choice of *g*^+^, when influence over outcomes is diminished. These plots conflate all values of rewards associated with *g*^+^, suggesting that the effect we observe is more to do with probability, rather than reward size. The effect is slightly stronger in low influence conditions.

In high influence blocks, I-scores positively correlated with the probability of success, obtained via simulations, of the choices made (*r* = 0.38, *p* = .02 95%CIs [.08, .72]), and inversely correlated with the average amount of money won when a win occurred (*r* = − .37, *p* = .03, 95%CIs [-.04, -.67]). Low I-scorers could sometimes attain *g*^+^ goals (which they more often pursued), making up for their frequent losses with rare but conspicuous wins. Thus, the effects of probability and winnings balanced each other out so that we found no significant relationship between the amount of money earned and I-scores in high influence conditions. In low influence blocks, we did not observe a correlation between I-scores and probability of success (*r* = 0.19, *p* = .24 95%CIs [-0.15, .49]); this is possibly because in low influence blocks the margin to make safer decisions is drastically reduced: all probabilities of success for all options are lower.

Although *g*^+^ choice proportion was significantly reduced when vehicles were less guidable (permutation test *p <* 0.001; both influence conditions), I- and C-scores were not associated with preference for the more guidable vehicle when choosing to attempt *g*^+^ (*p >* .14, I and C-scores; both influence conditions), or with the performance in discerning and choosing the most guidable vehicle on *Catch* trials (*p >* .11; I and C-scores, both influence conditions). This suggests, on a model-agnostic perspective (though this will be examined in more detail using modeling), that the microscopic action-to-outcome noise (or more abstractly, the objective quality of the tool used to carry out the plan) is not a particularly prominent feature of controllability in this task.

Finally, *g*^+^ choice proportion was strongly correlated across influence conditions (*r* = 0.65, *p <* .001, 95%CIs [.40, .81]), albeit being reduced when influence was low (permutation test *p <* 0.001). This suggests that the influence manipulation, and the drastic change in the overall achievability that follows (almost no chance of ever making *g*^+^ using any vehicle), only caused a rescaling, not a flattening, of proportionate choice. I and C-scores were not associated with differential proportions of *g*^+^ attempts across influence conditions, and so were not predictive of subjects’ choices’ sensitivity to the influence manipulation (I-scores: *r* = −.26, *p* = .13, 95%CIs [-.55, .07], C-scores: *r* = .25, *p* = .13, 95%CIs [-.08, .55]). Thus, perceived control is associated with behavioural signatures *within* each influence condition, but not with changes in strategy *across* the different influence conditions, despite the rather large difference in overall achievability between these.

#### 3.1.3. Single goal trials

I-scores correlated with choice of the closer vehicle in high influence blocks (*r* = .39, *p* = .025, 95%CIs [.06, .63]), but not significantly in low influence blocks (*r* = .26, *p* = .12, 95%CIs [-.07, .55]), regardless of the difference in guidability of the vehicles involved. No significant relationship was found for C-scores (*p >* 0.43 in either influence condition). I and C-scores did not predict rewards collected in SGTs, in either high influence (both *p >* .58), or low influence blocks (both *p >* 0.31); the latter is consistent with the fact that goals were only attained on a total of three occasions across all subjects and trials.

The frequencies of choices of the closer vehicle were strongly correlated across high- and low-influence blocks (*r* = 0.46, *p* = .005, 95%CIs [.15, .69]), reflecting similar decision-making strategies across influence conditions. We note that this result, in itself, constitutes evidence against a pure attainment probability-based class of models, as subjects had almost no chance of succeeding in low influence trials. As we shall see in our model-dependent analyses, this is also reflected in the poor performance of a pure probability-based model.

### 3.2. Model-dependent analyses

We complemented our model-agnostic descriptions of the data with a fully-fledged computational picture that provides a richer analysis of the sensitivities to trial features (distance, vehicular controllability and reward size), and more critically, probes the learning mechanisms driving choices of the riskier *g*^+^ goals. As anticipated in the methods, we employed a hierarchical graphical structure for all our models, in which sensitivities to trial features interacted additively to produce the value of choosing a vehicle-goal pair. We assume that subjects’ choices are algorithmically uniform across influence conditions (i.e., involve the same model components), but allow different sets of parameters for each condition. We generally report on our best computational accounts of both single and double goal trials and provide more detail about model construction and fits (e.g. recovered parameter distributions) in Supplementary Materials.

#### 3.2.1. Double goal trials

In the best model, subjects learned about (the odds of achieving *g*^+^ with) the two vehicles in a model-free way, throughout the task, using Rescorla-Wagner updates for their values, with separate learning rates for successes and failures. Vehicles then drive decisions in two ways: one that arises from the objective learning about their guidability (simply the update of a summary statistic of how many presses result in the correct movement), and the history of achievement which resulted from their use, in which successes and failures weigh differently, and in which the particulars of the problem at hand are absent.

The history of achievement of both vehicles is encapsulated in the terms *H*_*t*_(*v*_1_) and *H*_*t*_(*v*_2_). We show the evolution of the average of these terms (i.e. simply *H*_*t*_(*v*_1_)*/*2 + *H*_*t*_(*v*_2_)*/*2) through trials in figure 6, for each influence condition (A1, gray dots: high influence; A2, blue dots: low infuence). *H*_*t*_ is an experience-derived proxy for the probability of reaching *g*^+^ with a certain vehicle. This term (which takes values in the interval [−1, 1]; −1 meaning lowest, and +1 highest, perceived achievability) evolves through Rescorla-Wagner updates: it increases when a *g*^+^ goal is achieved, and it decreases when *any* loss is incurred. For vehicle *v*_1_ (the same exact updates apply, of course, to *v*_2_), this will evolve as per the equation:

**Figure 6:**
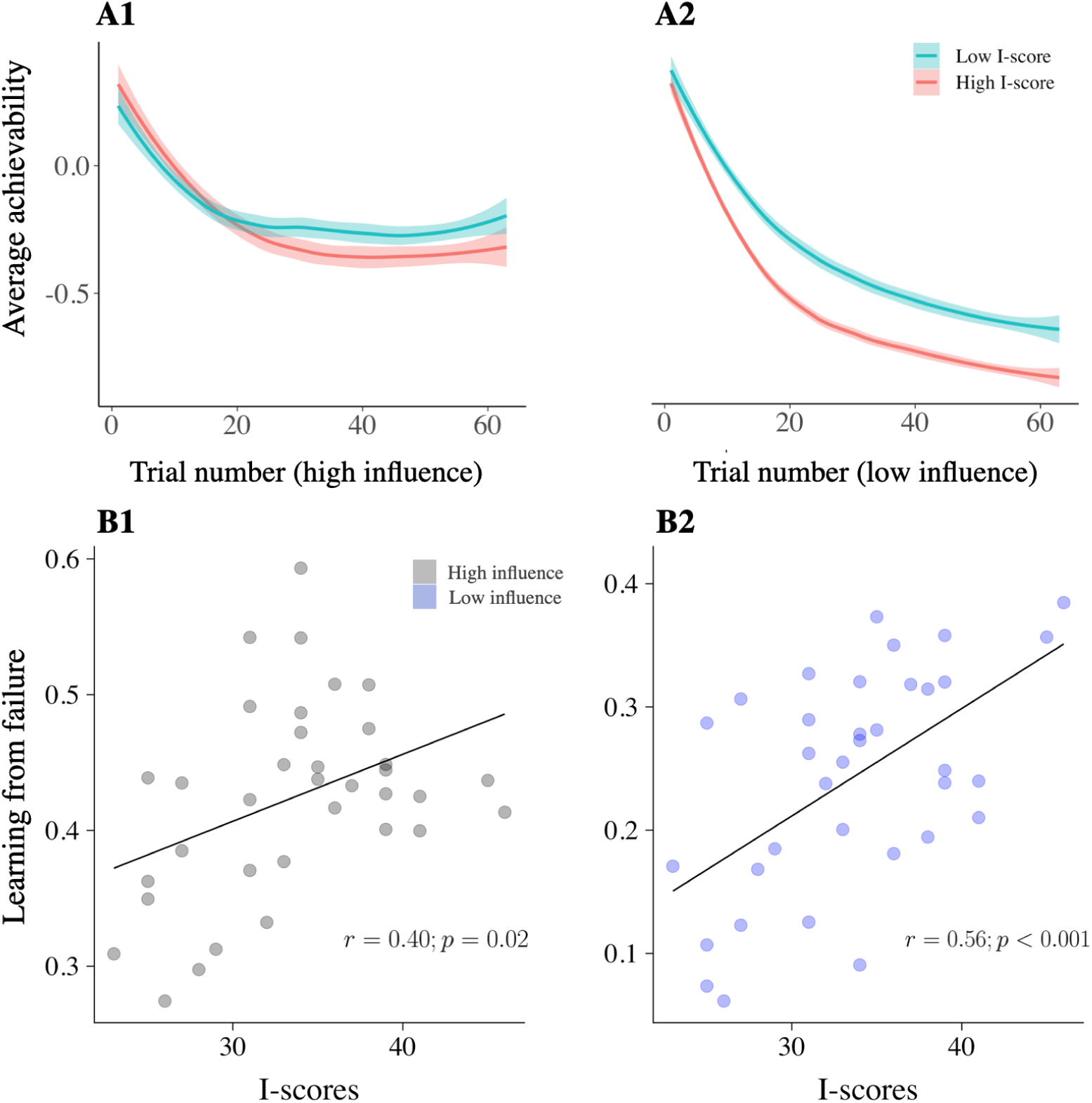
Plots **A1** and **A2** illustrate the evolution of average achievability (i.e. the parameter *H*_*t*_) through DGTs in high and low influence conditions respectively. This is the *average* achievability because we consider the average of the values of *H*_*t*_ for the two vehicles at each trial. The shaded error bands reflect standard error across subjects. The *x*-axis reports the ordered trials for each condition. This is a total of 64 trials (as there are 16, times 4, DGTs per each condition). For illustration purposes, we divided our subjects into two groups using a median split on I-scores. We can observe the average achievability learnt through Rescorla-Wagner updates decrease faster for high I-scorers in both influence conditions, however, the difference across the two groups is most apparent in low influence conditions. Plots **B1** and **B2** illustrate the correlations between I-scores and learning from failure in high and low influence conditions respectively, which underpin the temporal evolution of *H*_*t*_ shown in A1 and A2. The effect is stronger in low influence conditions.

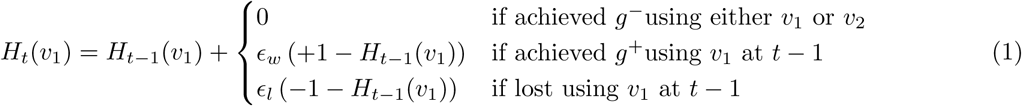

Here, *t* ∈ {1 … 16} indicates the current trial. When *t* = 1 (first trial of the task), *H*_1_ is equal to the prior value for achievability. *ϵ*_*w*_ (*ϵ*_*l*_) are the learning rates for success (failure). Note that this update only takes place for the vehicle that was chosen at time *t* − 1, in this case *v*_1_. The value of *H*_*t*_ for *v*_2_ remains the same, i.e. *H*_*t*_(*v*_2_) = *H*_*t*−1_(*v*_2_). The rationale of this learning rule is that learning by experience in this task should take into account that (1) failure to achieve *any* goal should imply that the odds of achieving *g*^+^ should be lower (as *g*^+^ is always harder); (2) success in attaining *g*^−^ should not rationally imply that *g*^+^ is more achievable (since *g*^−^ goals are always easier) (3) achieving *g*^+^ is the only evidence that counts towards achievability of *g*^+^ with a certain vehicle. For completeness, we tested a further model which had a separate learning rate for achievements of *g*^−^ goals, which performed less competently.

In table 2, we report the out-of-sample performance for all our models. In Supplementary Materials, we show the general calibration plots for choice frequency of the more rewarding goals (*g*^+^) and more controllable vehicles (*v*_1_), demonstrating the generative adequacy of the winning model.

**Table 2:** Model comparison for DGTs. We report out of sample likelihoods for all our models. Starting from the easiest (additive), which forms the basis for all our models, to the most complex (and best performing). Pairwise comparisons between individual trials’ out-of-sample likelihoods of model 6 against all other models yielded *p* values *<* 0.001.

| Models | LOSO |
| --- | --- |
| Additive (1) | 0.623 |
| $g^+$ bias (2) | 0.632 |
| Temporally evolv. bias (4) | 0.602 |
| Win-stay Lose-shift (3) | 0.644 |
| Vehicle indep. R-W (5) | 0.637 |
| Vehicle dep. R-W (6) | 0.653 |

Post-hoc correlations of the parameters from the winning model revealed that the main determiners of the relationship between reduced choice of *g*^+^ and I-scores were the increased learning rates (learning from failure) of the model-free term (in both high, and low, influence conditions) and increased distance sensitivities (in high influence conditions). In low influence conditions, we registered lower priors for achievability in high I-scorers. These low levels are likely indicative of fast learning of inachievability post introductory trials in low influence conditions.

Reward sensitivities anti-correlated significantly with I-scores, but they were not essential for the recovery of the main effect of reduced *g*^+^ choice up to permutations when compared to other variables (see figure 7). Finally, in order to obtain a global view of variable importance (not towards the recovery of the model agnostic effect concerning choice frequency of *g*^+^, but towards prediction of I-scores) we used random forests (‘party’ package in R) to predict I-scores using all subject-wise parameter estimates. Here, variable importance was measured as the mean increase in the squared residuals of predictions (estimated with out-of-bag cross-validation) as a result of mean parameter estimates being randomly shuffled. The results were consistent with what we found for the recovery of the main model-agnostic effect, but placed an even stronger emphasis on the importance of learning rates because of their relevance to the prediction of I-scores (see figure 7).

**Figure 7:**
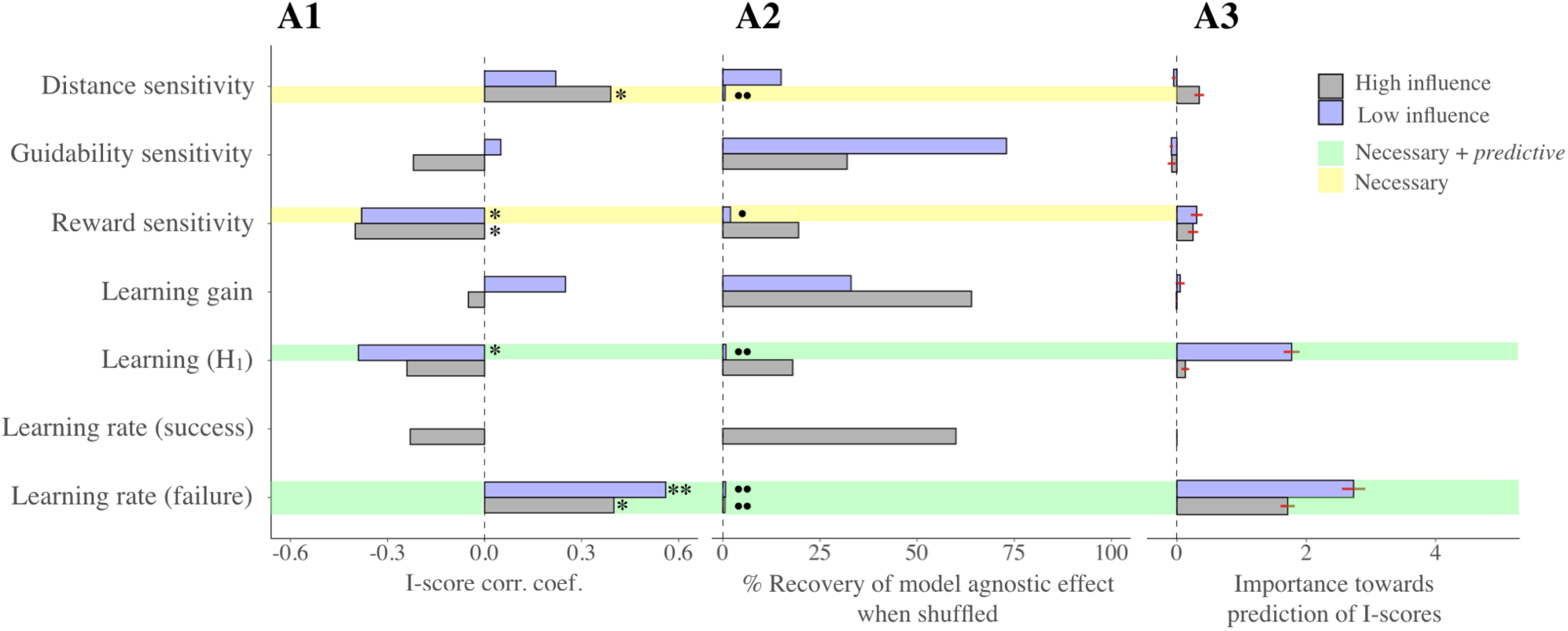
I-scores and parameter estimates. This series of plots illustrates the relationship between I-scores and the recovered parameter estimates of the winning model (i.e. ‘Vehicle dep. R-W’, in table 2) in three ways. Note that learning rate from success (in attaining *g*^+^) is absent in low influence conditions, as only 3 wins over all datasets were recorded. **A1** illustrates the correlation coefficients between each parameter and I-scores. Next to each bar, a star (*) indicates significant correlations post false discovery rate correction for multiple comparisons. Two stars (**) indicate significant correlations post Bonferroni correction for multiple comparisons. In **A2**, each bar exemplifies the percentage of recovery of an equal or larger correlation coefficient of I-scores and *g*^+^ choice frequency (the effect found in model-agnostic analyses) when shuffling the recovered parameters across subjects. We performed this analysis (described in methods) to understand which parameters played the most important role in driving this effect. Two circles next to the bar indicate a percentage of recovery smaller than 1%, one a percentage smaller than 5%. Thus, according to this measure, the effect is driven by distance sensitivities (the most apparent feature of achievability) in high influence conditions, and (in both influence conditions) by the learning rates from failure. Finally, in **A3**, each bar shows the loss in predictive power of I-scores when using the recovered parameter estimates as predictors. This measure used random forests, and was obtained as described in Methods. The errorbars signify the standard errors (the procedure was repeated 50 times). Finally, as the legend shows, the parameters highlighted in green are those which show significant effects with I-scores, are needed to reproduce the model agnostic effect, and exhibit a very high importance for I-score prediction. In yellow, we highlighted those parameters which, albeit being necessary to reproduce the model-agnostic effect, turned out not to be critical for predicting I-scores.

#### 3.2.2. Single goal trials

In table 3, we report results for all our models for SGTs. We found that a model based on the simple additive interaction of vehicular guidability (which, by the time of the SGTs, we assume to be known), vehicle-goal distance, and their interaction, yielded the most parsimonious account of the data. Consistent with our model agnostic findings, we found no significant relationship for C-scores, while I-scores correlated with distance sensitivity in high influence blocks (*r* = .40, *p* = .01, 95%CIs [0.08,0.65]). Internals then appealed to a more apparent cue of achievability (the vehicle-goal distance) in order to maximise the probability of success in SGTs; however, this did not quite work out in the same way as with DGTs, as here I-scorers did not make more probably successful decisions on average. By design, in fact, it is best to choose the central vehicle in two out of the four SGTs at the end of each block (this is described in figure 2).

**Table 3:**
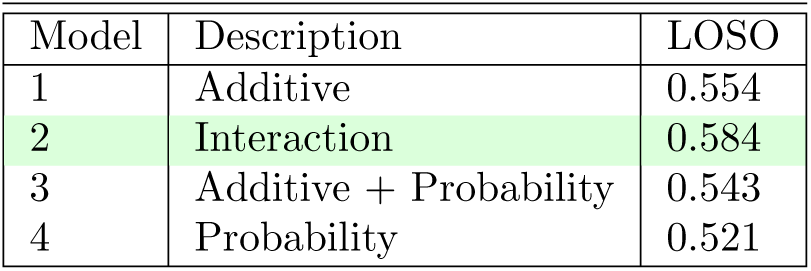
Model Comparison for SGTs. The most parsimonious model accounts for data through adding distance and vehicular guidability features, and their multiplicative interaction. I-scores were explicative of distance sensitivities, but only in high influence blocks.

| Model | Description | LOSO |
| --- | --- | --- |
| 1 | Additive | 0.554 |
| 2 | Interaction | 0.584 |
| 3 | Additive + Probability | 0.543 |
| 4 | Probability | 0.521 |

Importantly, the focus on distance in SGTs reflects the value attributed to this feature during DGTs; distance sensitivity parameters were in fact correlated across S and DGTs (high influence: *r* = 0.49, *p* = 0.002, 95%CIs [.19, .70]; low influence: *r* = 0.65, *p <* 0.001, 95%CIs [.35, .78]).

Perhaps due to the near flat probability of reaching the goal inherent of low influence conditions SGTs, neither distance nor vehicular control sensitivities correlated with I-scores (distance s.ty: *r* = 0.21, *p* = 0.22, 95%CIs [-.13, .5]; veh. guidability s.ty: *r* = −0.30, *p* = 0.08, 95%CIs [-.57, .03]).

### 3.3. Exploratory analyses

#### 3.3.1. Powerful others

We did not have preliminary hypotheses regarding the P-scale. However, additional analyses of our data revealed that money earned in low influence DGTs (but not SGTs, recall these almost always led to losses by design) was very strongly positively correlated with the P scale *r* = .57, *p <* 0.001, 95%CIs [.3, .76]); P-scorers’ choices in low influence trials were also less likely to lead to a loss, whichever the amount (*r* = − .49, *p* = 0.003, 95%CIs [.18, .7]). See figure 8. We investigated which parameters could be responsible, rather as we did for I-scores, and found that the distance sensitivities parameters correlated heavily with P-scores (*r* = .48, *p* = 0.003, 95%CIs [.17, .7]) and were necessary to recover the effect of increased earnings (*p* = 0.02) and choice probability.

**Figure 8:**
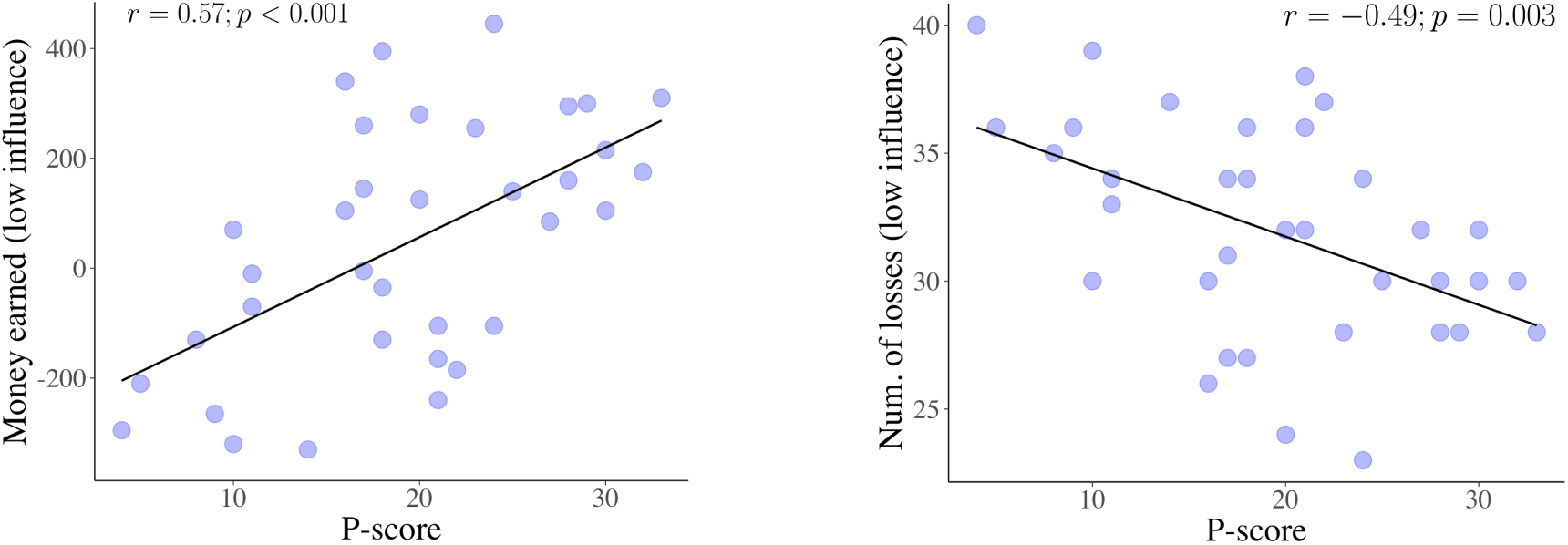
Plots illustrating the positive correlation between P-scores and money earned (left), and the negative correlation between P-scores and number of losses (right), in low influence trials.

## 4. Discussion

We designed a novel decision-making task which requires subjects to learn and adapt to the controllability of environments, and thereby allows us to isolate distinguishable components relevant to control. We analyzed the results using both model-agnostic assessments of choice frequencies and model-dependent characterizations of the trial-by-trial effects of rewards and punishments.

We found correlations between specific locus of control (LoC) scores and both model-agnostic and model-dependent aspects of performance and learning. Internal (I)-scores correlated with a form of “timidity” in the shape of reduced choice of the less attainable, more rewarding (*g*^+^) goals. This correlation was significant when subjects had either greater or lesser influence over the environment, but was stronger in the latter condition. In the high influence condition, the timidity of high I-scorers entailed that they had a higher likelihood of success (albeit not gaining more points on average, because they chose the impoverished goals).

Our computational analyses favoured a model in which value-based decision making was based on a form of Rescorla-Wagner learning of how achievable the more attractive goal (*g*^+^) was with each vehicle (a term we call *H*_*t*_). There was evidence for separate learning rates for appetitive (success) and aversive outcomes (failure)(Shapiro et al., 2001; Frank et al., 2004, 2007; Niv et al., 2012; Daw et al., 2006; Palminteri & Pessiglione, 2017), with the latter being higher, as has previously been observed in other circumstances (Niv et al., 2012; Gershman, 2015).

We then showed that the boosted learning from failure and biased initialisation of achievability (*H*_1_) were instruments of the negative relationship between *g*^+^ goal choice frequency and I-scores observed in our model-agnostic analyses. Crucially, our random forests-based analyses established that these were substantially the best predictors of internality.

Our observations are broadly consistent with existing observations about the role of LoC in decision making, including the fact that internal subjects tend to place more conservative bets (Liverant & Scodel, 1960), opt for courses of action allowing lower chances to incur in episodes of failure (Julian et al., 1968), and expend more effort in a task when its goal is framed as avoiding punishment, rather than attaining reward (Gregory, 1978). However, our findings broaden these results through our focus on learning.

Closer to our focus is the work of Rotter and his group, who utilised explicit *verbal* manipulations of the contingency between actions and outcomes to study the evolution of explicit expectations (Rotter, 1966). Different subject groups would typically be provided with different sets of instructions, with the same task framed as a matter of “Skill”, or “Chance” (there would sometimes also be “ambiguous” instructions). The impact of reinforcement history on expectations over future outcomes was then probed.

Changes in expectation were blunted in “Chance” conditions (Rotter, 1966; Phares, 1957; James & Rotter, 1958; Rotter et al., 1961). Under “Skill” conditions, subjects quickly increased their expectations of success after wins, and only slowly learned their inability to succeed after failure (Rotter et al., 1961). The discrepancy between this result and our observation of faster learning from failure might arise from a difference between control (as in I-scores) and the spectrum between skill and chance (as in Rotter’s work). That we found the LoC ‘Chance’ C-score to be unrelated to any aspect of performance suggests that the task was largely perceived as a matter of skill. This, in turn, makes it very unlikely that the learning effect we observe is due to a form of gambler’s fallacy (i.e., an expectation of achieving *g*^+^ now because it has not been achieved in a while), as this latter is more of a feat of high beliefs in Chance (Rotter, 1966).

By contrast, the strong link between the LoC ‘Powerful-Other’ P-scores and performance in low influence conditions was unexpected, and so merits replication. One possibility is that subjects with high P-scores might be more fatalistically adaptive to circumstances in which a source of control exists, but is beyond reach. It is worth noting that the experimenter was present throughout the experiment, possibly increasing discomfort for those with low P-scores, and affecting performance.

The tie we observed between high I-scores and boosted learning from failure is relevant to converging evidence from multiple different disciplines about the role of perceived controllability in coping in the face of negative events (e.g., Lefcourt, 1976; Anderson, 1977; Lefcourt, 2013), or stress (Abouserie, 1994; Sandler & Lakey, 1982). Perceived control regulates emotional and behavioural responses to aversive outcomes, both when availing of situational manipulations (see for instance Maier & Seligman, 2016), or using LoC questionnaires to measure generalised expectations (Hiroto, 1974; Lefcourt, 1976; Harnett et al., 2015). For instance, perceiving an aversive stimulus as controllable modulates the neural responses associated with its presentation (Salomons et al., 2004) or its predictive cue (Wood et al., 2015), and reduces anticipatory anxiety (Kerr et al., 2012). The picture that emerges from combining this literature with our results is one where dysfunctional reactions to perceived unachievability (excessive frustration, stress, or anticipatory anxiety; such as those found in Julian et al. (1968)’s second experiment) might be behaviourally underpinned by a blunted capacity to learn from (and adapt to) it. To make matters worse, this mechanism easily self-sustains: failure without adaptation entails more exposure to failure, and is bound to reinforce an idea of low internality.

In neural terms, Harnett et al. (2015) found that learning-related changes in the emotional response to negative outcomes were mediated by activity in ventromedial prefrontal cortex (vmPFC); crucially, as individuals moved towards the external end of the LoC spectrum, the vmPFC response to predictable threat decreased (the study, however, could not differentiate between internality, chance and powerful others). Our findings provide a possible instrumental amplification of these results. If LoC internality is linked to aversive learning rates, as our results show, it is possible that a generalised idea (or prior) on internality might moderate learning rates from aversive events through vmPFC. At the same time, recent work implicates the vmPFC in an optimism bias effect (Kuzmanovic et al., 2018). The authors found that vmPFC activity was directly associated with the integration of positive information (i.e. it increased with retaining of good news and decreased retaining of bad news; the information was, for instance, about subjective perceived risk of developing an illness, expressed as a percentage). This is in apparent contradiction with our suggestion that vmPFC might be implicated in faster adaptation to failure (and therefore negative information). But this is only apparent, since, if vmPFC is broadly implicated in a more “proactive” outlook on the environment, protecting us from courses of action which will make matters worse, then it makes sense that it might skew the statistics positively, positing that there is something we can do to change those statistics for the better. Finally, the vmPFC has duly been implicated in computations involving control detection (Kerr et al., 2012; Salomons et al., 2004; Wood et al., 2015; Harnett et al., 2015; Christianson et al., 2009; Wang, 2019), and, in studies in rats, has been shown to suppress the over-exuberant activity of serotonergic neurons in the dorsal raphe when they have control over aversive outcomes (Maier & Watkins, 2005). The latter is one of the sources of evidence that serotonin is involved in aversive learning (Daw et al., 2002; Boureau & Dayan, 2011; Dayan & Huys, 2009), and could be involved in the excess learning from failure that we found.

Despite these promising findings, we should note some caveats which concern our results. For instance, while we found no significant relationships involving the Chance sub-scale, it is entirely possible that these might exist, but simply play less apparent roles than the I- (or P-) sub-scales. Replications of our study, possibly availing of larger sample sizes, could offer further support to our results. From a design perspective, it would be desirable to have a range of different losses, so that sensitivity to negative outcomes could be assessed; it would also be important to present the threat of primary aversive outcomes such as shocks. Further, we implemented control in a rather direct manner, using effort. It would be interesting to see if our findings generalized to a simpler choice task. Finally, the task lacked the sort of complexity that would allow a distinction between different mechanisms for choice such as model-based and model-free learning and planning (see, e.g. Gläscher et al., 2010). Controllability, and indeed biased learning from success and failure, might affect each differently.

In sum, we have provided a new tool for investigating controllability, and new results about the link between internality and biased aspects of learning. Our findings are consistent with the proposition that internality is subservient to giving negative information more weight, in a way that facilitates adaptation, or as Lefcourt would put it, more positively (Lefcourt, 1976).

