## Supplemental Materials for "Subjective Beliefs In, Out, and About Control: A Quantitative Analysis"

### Supplementary Materials

#### Example of block structure (DGTs)

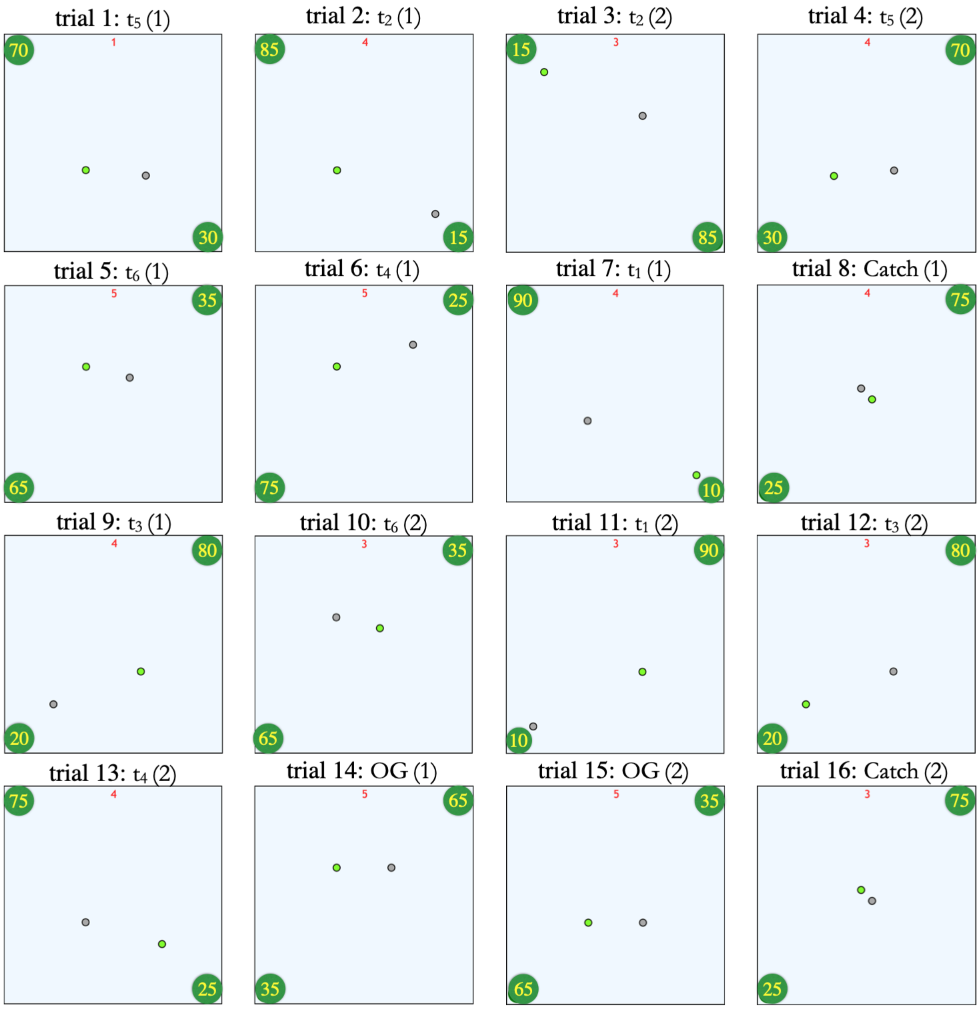


Figure 1: **Instance of a block from trials 1 to 16 (DGTs)**. The background of the canvas is light blue, indicating the influence condition (for half of the subjects this would signify high, and for the other half, low, influence). The vehicles and goals for the first 16 trials of a block are displayed as they appear at the start of the trial (vehicle selection phase; the red timer on top denotes time left to choose, random here) with goals made slightly bigger, for illustrative purposes. The ordering of the trials is randomized, as is the position of the goals (the angle by which they are rotated). The labels for each trial type are on top of each screenshot, the number in parentheses being the order of occurrence of this trial type within the block. Note how on the second occurrence (i.e. (2)), the vehicles position is swapped. The sixteenth trial (bottom-right) is *always* a catch trial; but the ordering of all the other trials is random.

#### Model-dependent analyses

**Double goal trials** We first posited that subjects simply choose according to their preferences for achievability and reward. Thus, our basic modelling approach combined, additively, the main trial features driving choice, i.e. distance, guidability and reward. We then incrementally (using model complexity control) built more sophisticated, and better performing, models, to uncover temporal dependencies across trials (learning).

In its most basic form, $U_t(v, g)$, the utility of choosing vehicle $v$ to attain goal $g$ at trial $t$, is modelled as follows:

$$U_t(v, g) = -\alpha_d \cdot d_t(v, g) + \alpha_v \cdot 100\gamma_t(v) + \alpha_r \cdot r_t(g) \quad (1)$$

The expression considers the values, at trial $t$, of the euclidean distance between vehicle $v$ and goal $g$ (i.e. $d_t(v, g)$; which is obtained by dividing the Manhattan distance by 2 and multiplying by $\sqrt{2}$), the guidability of $v$ as inferred by trial $t$ (i.e. $\gamma_t(v)$; this is multiplied by 100, so that the guidability values lie in the same range as distances and rewards, and parameter estimates are comparable), and of the reward connected to $g$, i.e. $r_t(g)$. There are, of course, four possible choices of vehicle-goal pairs in all DGTs. A subject could in fact choose either vehicle ($v_1$ or $v_2$) and aim for either goal ($g^-$ or $g^+$). The probability of each option is modelled as a generalization of the softmax function for four options (an ordinary multinomial logistic regression):

$$P(v, g) = \frac{e^{U_t(v,g)}}{\sum_{v' \in \{v_1, v_2\}, g' \in \{g^-, g^+\}} e^{U_t(v', g')}} \quad (2)$$

This is our additive model for DGTs ("Additive"), and constitutes the starting structure of *all* our models. In turn, these differ according to the ways they refine $U_t(v, g)$, adding extra features.

In the task, distances and rewards are directly observed. However, vehicle guidability (i.e. $\gamma_t(v)$) has to be learned. We consider a Bayesian form of learning for both vehicles, as follows:

$$\gamma_t(v) = \frac{4}{3}R - \frac{1}{3}$$
$$\text{where } R = \frac{\sum_{\tau=1}^{t-1} n_{good}(\tau)}{\sum_{\tau=1}^{t-1} n_{good}(\tau) + \sum_{\tau=1}^{t-1} n_{bad}(\tau)} \quad (3)$$

$n_{\{good,bad\}}(t)$ gives the number of {good,bad} moves at trial $t$. $R$ is the perceived probability that the vehicle will go in exactly the direction we press towards. Note that this can arise both as a consequence of the vehicle actually following our command, i.e. $\gamma_t(v)$, or because the vehicle *randomly* went in the direction we asked it to, i.e. $\frac{1}{4}(1 - \gamma_t(v))$. So to infer $\gamma_t(v)$ from the statistic $R$ we need to solve $R = \gamma_t(v) + \frac{1}{4}(1 - \gamma_t(v))$ , which yields the first line of the equation. At trial 1, when either vehicle is fully unknown, we set $n_{good} = 1$ and $n_{bad} = 1$, yielding a controllability $\gamma_1(v) = 0.33$. We also explored the possibility of drawing these initial values from top level priors, but ultimately kept them fixed, for simplicity.

In sum, our basic model simply treats additively the set of factors driving choice. At first glance, one might argue that since distance and reward are anti-correlated by design, inference will face an unidentifiability issue, as sensitivities to distance and reward are confounded (increasing sensitivity to reward or diminishing that to distance would yield approximately the same log-likelihood); however, trial type $OG$ (see e.g. figure 1), which appears twice in DGTs, was conceived to resolve exactly this issue, as it makes the larger goal closer to both vehicles, thereby breaking the anti-correlation.

##### Learning of achievability in DGTs

We added extra model components to account for learning of achievability throughout the task, reasoning that we might well observe aspects of model-free learning, on top of value-based choice. We then decided to keep the additive structure of equation (1) and add a time-dependent component.

**Winning model ("Vehicle-dependent RW").** In DGTs, in all but 'Catch' trials, one vehicle ($v_{c_t}$) is at the center of the canvas, while the other is closer to the less rewarding goal ($g^-$). We then hypothesized that the main form of learning would be about the success in achieving $g^+$ using the central vehicle, as subjects are faced with the recurrent question whether $g^+$ will be achievable with $v_{c_t}$ (empirically, most attempts to $g^+$ took place utilising the central vehicle, i.e. 92%). Recall that the distance (36 Manhattan steps, or $18\sqrt{2}$ euclidean) between the central vehicle and $g^+$ was always the same, so the main learning factor underlying choice of $g^+$ could only be connected to the the vehicle itself. This form of achievability is essentially model-free, and only depends on the experience gathered with the central vehicle.

To include a learning term, and test whether its use is justified, we kept all the "static" aspects of decision making as per the basic structure in equation 1, simply adding an extra component $H_t(v_{c_t})$ to the expression for $U_t(v_{c_t}, g^+)$, leaving unchanged the remaining three terms:

$$U_t(v_{c_t}, g^+) = -\,\alpha_d \cdot d_t(v_{c_t}, g^+) + \alpha_v \cdot \gamma_t(v_{c_t}) + \alpha_r \cdot r_t(g^+) + \alpha_H H_t(v_{c_t}) \tag{4}$$

where $H_t(v_{c_t}) \in [-1, +1]$ is the output of a learning rule (see below) that integrates the history of successes and failure, and $\alpha_H$ is a parameter ("learning gain") that quantifies the effect of this term. We should also point out that $d_t(v_{c_t}, g^+)$ is simply equal to $18\sqrt{2}$, a constant, since the distance between the central vehicle and $g^+$ is fixed throughout.

Of course, many learning processes could be suitable, so there is a question as to how $H_t$ should be updated given success and failure. The simplest possible learning rule is the Rescorla-Wagner (RW; delta rule) update, according to which subjects start out on the initial trial of the task with an initial guess for the achievability of $g^+$ from the center, $H_t(v_{c_t})$ (i.e. $H_1$). This would then evolve on a trial-by-trial basis. On each trial, there are 8 scenarios that this learning rule should consider: a subject could choose either vehicle, aim for either goal, and face either of two fates.

In the winning model, the notion of success and failure which affect $H_t(v_c)$ is as follows. We regard, as success, only that in gaining the $g^+$ goal (regardless of whether the central or the other vehicle was chosen), and as failure, any failure both in an attempt to a $g^-$ or $g^+$ goal. This is because $g^-$ is invariably easier to reach than $g^+$, so a loss when attempting the easier goal should rationally imply that $g^+$ is less achievable than we thought. Conversely, succeeding in attaining $g^-$ does not imply that $g^+$ will be more achievable. Note that since we only count successes on $g^+$ goals towards the update of $H_t(v)$, learning from wins is not meaningful in low influence blocks, where all (but three attempts across all datasets in total) to $g^+$ led to a loss. Thus, as one would expect, learning of the hopeless achievability of $g^+$ in low influence conditions only happens through failures. We have of course tried variations of this model which took into account different learning speeds for wins and losses faced when attempting $g^-$ and $g^+$. While we do not report all of these here, for brevity, their performance was not as competent as our final formulation.

Thus, in the winning model, the evolution of $H_t(v)$ across trials is:

$$H_t(v) = H_{t-1}(v) + \begin{cases} 0 & \text{if achieved } g^- \text{ using } v \text{ at } t-1 \\ \epsilon_w \left(+1 - H_{t-1}(v)\right) & \text{if achieved } g^+ \text{ using } v \text{ at } t-1 \\ \epsilon_l \left(-1 - H_{t-1}(v)\right) & \text{if lost using } v \text{ at } t-1 \end{cases} \tag{5}$$

where $t \in \{1 \dots 16\}$ indicates the current trial, and $v$ identifies the vehicle that was chosen at trial $t-1$. For the vehicle that was *not* chosen at trial $t-1$ the value for $H_t$ is simply copied over, i.e. if we denote this vehicle as $\bar{v}$, then we have $H_t(\bar{v}) = H_{t-1}(\bar{v})$. $\epsilon_w$ ($\epsilon_l$) are the learning rates for success (failure). This expression keeps track of a surrogate value for the probability of achieving $g^+$. As such, it does not distinguish between the amount won or loss in the previous trial.

As we model learning throughout the task, at the end of each block in high/low influence conditions, the achievability value for each of the two vehicles involved, i.e. $H_t(v)$ is averaged and used as the starting value for the next block of the same influence conditions.

**Other models.** Prior to testing our final formulation, we tested several simpler learning schemes. Aside, of course, from our basic additive model (equation 1) we tested four other ways to refine this, which we describe below. These models' out of sample performances as measured by average leave-one-subject-out likelihoods are reported in the main text.

- *Bias.* The bias model is simply the basic structure of equation 1, endowed with a free parameter to account for a propensity to choose $g^+$ that is not dependent on any other characteristic of the trials. Thus, it simply adds a term to only the utility of attaining $g^+$ with the central vehicle:

$$U_t(v_{c_t}, g^+) = \alpha_d \cdot d_t(v_{c_t}, g) + \alpha_v \cdot \gamma_t(v_{c_t}) + \alpha_r \cdot r_t(g) + \beta_{bias} \tag{6}$$

  The term in question is of course $\beta_{bias}$.

- *Temporally evolving bias.* This model allows for temporal evolution of choice throughout the task, but does so without considering information about success and failure. Thus, we have here the same formulation as in equation 4, yet $H_t(v_{c_t})$ becomes a term $H_t$, where:

$$H_t = H_{t-1} + \alpha\left(+1 - H_{t-1}\right) + (1-\alpha)\left(-1 - H_{t-1}\right) \tag{7}$$

  Note that there is no dependency of $H_t$ on vehicles. This is simply a version of the *Bias* model where bias is time varying, as described by $\alpha$, rather than constant. The subjects' tendency to see $g^+$ as more (less) achievable as the task unfolds depends on $\alpha$ being greater (lower) than 0.5. Testing this model allows us to discern whether the tendency to choose $g^+$ from center depends on the history of success and failure, or on some task-independent feature (e.g. progressive disengagement from the task, or fatigue).

- *WSLS*. The Win-stay Lose-shift formulation only has a 1-trial memory to (de)incentivate choice of $g^+$ with a vehicle. In this case $H_t(v_{c_t})$ evolves as:

$$H_t(v_{c_t}) = \begin{cases} \epsilon_w & \text{if achieved } g^+ \text{ using } v_{c_t} \text{ at } t-1 \\ \epsilon_l & \text{if lost using } v_{c_t} \text{ at } t-1 \end{cases} \tag{8}$$

  Of course, $\epsilon_w > 0$ and $\epsilon_l < 0$.

- *Vehicle-independent RW*. The last alterantive, which is perhaps closest to the winning model is a Rescorla-Wagner rule that does *not* keep separate records for the two vehicles available inside a block. Here, achieving $g^+$ with a vehicle will simply increase the value for choosing $g^+$ regardless of which vehicle is used. That this model performs worse than our winning model means that subjects must be using the different vehicles' histories of achievements to make their decisions, and are not simply being motivated by success and failure to try again (or quit trying) to achieve $g^+$. Here, the $H_t$ term drops the dependency on vehicles, so equation 5 becomes:

$$H_t = H_{t-1} + \begin{cases} \epsilon_w\,(+1 - H_{t-1}) & \text{if achieved } g^+ \text{ at } t-1 \\ \epsilon_l\,(-1 - H_{t-1}) & \text{if lost at } t-1 \end{cases} \tag{9}$$

**Single goal trials.** In SGTs, the expression for $U$ (i.e. equation 1) omits the term concerning reward, since only one goal is available. Further, because of the large number of preceeding DGTs, we make the assumption that $\gamma_t(v)$ has fully converged to the true controllability value of the vehicle. Equation 1 then becomes:

$$U(v) = \alpha_d d_t(v) + \alpha_v 100\gamma(v) \tag{10}$$

here $\gamma(v)$ is the true value for the controllability of the vehicle, that is, one of 0.28,0.36,0.43. Our interaction model (the winning model for SGTs) posits a further term:

$$U(v) = \alpha_d d_t(v) + \alpha_v 100\gamma(v) + \alpha_{int}\gamma(v) \cdot d_t(v) \tag{11}$$

$\gamma(v) \cdot d_t(v)$ is an interaction term of guidability and distance.

Finally, we have the probability (P) model, which only used sheer probability as a predictor. Here, the expression for $U$ is simply $U(v) = \alpha_p \omega(\gamma(v); d; f)$. In this expression, $\omega$ denotes the probability of reaching the goal when choosing vehicle $v$ and pressing in an optimal manner. This term, whose approximation we also describe in SMs, depends on the distance from the goal ($d$), the controllability of the vehicle $\gamma(v)$, and the frequency of pressing that the subject expects to exert over the vehicle. We allowed for subjects' frequencies to vary, so that the $f$ that $\omega$ depends on is indeed a free parameter which depends on subject and condition. In high (low) influence conditions individual $f$ parameters were drawn from fixed, top-level normal distributions with mean 8 (4) and standard deviations of 1.

#### Details of modelling results

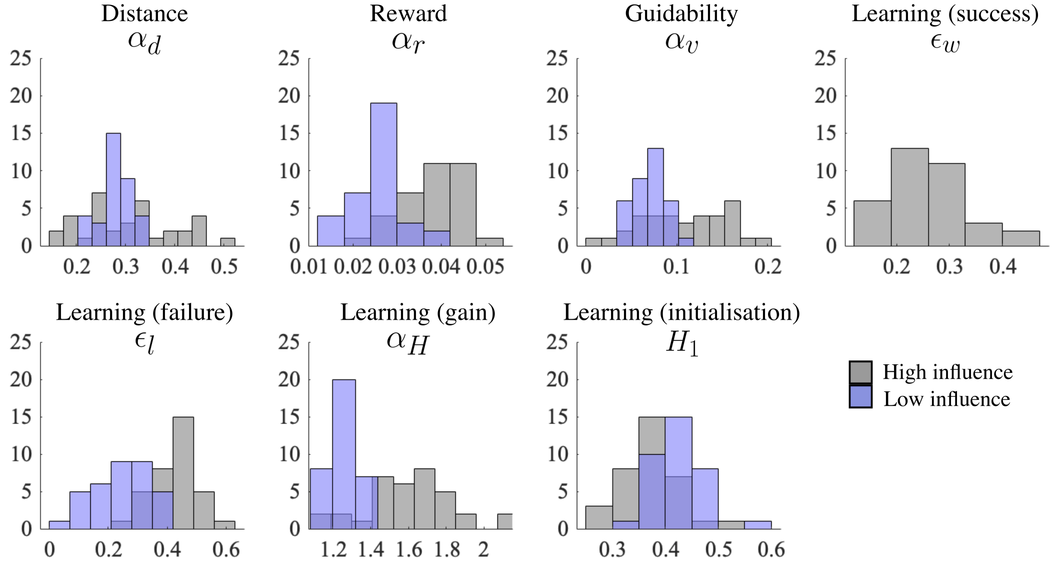


Figure 2: **Winning model ("Vehicle dep. RW"): recovered parameter distributions**. Here, we show the recovered parameter fits for our winning model, obtained via stan. Learning from success ($\epsilon_w$) is inherently only defined in high influence conditions. All values are on average lower in low influence conditions, possibly due to the greater stochasticity in decision making (this is equivalent to having a lower inverse temperature). $H_1$ values in low influence conditions are slightly higher, possibly compensating for the incapability of the model to describe higher, initial tendencies to choose the riskier goal through other trial-based features. Although learning from failure ($\epsilon_l$) parameter fits are lower on average in low compared to high influence conditions, recall that this is the only learning feedback we posit for low influence conditions. Thus, the resulting $H_t$ values will, in time, quickly fall below those reached in high influence conditions, as they can only monotonically decrease.

#### Trial-wise model comparisons

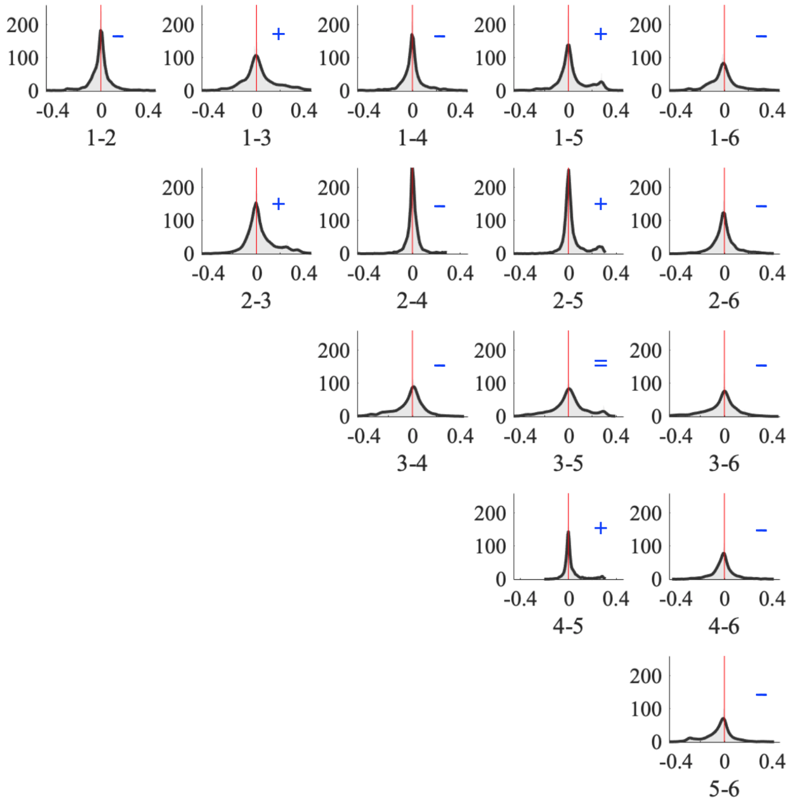


Figure 3: **Trial-wise, out-of-sample performance comparisons**. This matrix of smoothed histograms shows the distributions of differences in log-likelihoods when fitting the data to subjects as they are left out of training. The first row shows the difference across models 1 and all models through to 6 (i.e. 1-2 through to 1-6). The same applies to rows 2 to 5. All differences are strongly significant ($p < 0.001$, due to the high power of considering all trials for all subjects) - except for the non-significant difference between models 3 and 5 ($p = 0.10$), and the significant one between models 2 and 4 ($p = 0.01$). The models are reported with the same numbering as that in which they appear in the main text, that is: 1 = "Additive", 2 = "$g^+$ bias", 3 = "temp.evolv.bias", 4 = "Win stay, Lose shift", 5 = "Vehicle indep. RW", and 6 = "Vehicle dep. RW". The blue sign, top-right of each plot, reports the prevalent sign of the difference between the out-of-sample log-likelihoods. All models performed worse than n.6.

#### Account of summary data statistics in DGTs

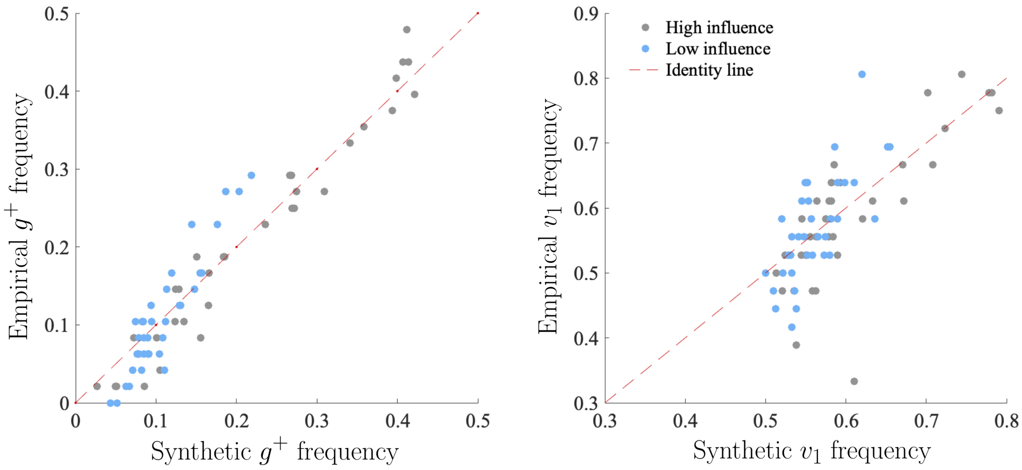


Figure 4: **Calibration of goal and vehicle choice frequencies**. These plots show that the winning model ("Vehicle dep. RW") generated frequencies of $g^+$ choices (left) and more controllable vehicle $v_1$ (right) are well calibrated to the data. Each dot is a subject's statistic as computed in a particular influence condition (gray: high; blue: low). The variance explained by the model is high for both measures ($g^+$ and $v_1$ choice frequency) and influence conditions (high and low): (i) $g^+$: high influence, $r^2 = 0.96$, low: $r^2 = 0.84$, (ii) $v_1$: high influence, $r^2 = 0.69$, low: $r^2 = 0.49$. Low influence statistics are less faithfully recapitulated than high influence statistics, possibly on account of subjects inherently attempting to randomise their decision making to obtain more success. Finally, vehicular choice is less faithfully recapitulated in both influence conditions – this is expected, since subjects only have a noisy notion of which vehicle is most controllable.

#### Computation of Probabilities

In our task, the probability of reaching a goal with a certain vehicle can be computed in two ways. In our model agnostic analyses, and when fine tuning the task (e.g. to choose the guidability levels of the vehicles), we used synthetically derived proportions of successful attempts. We obtain these by simulating episodes in which artificial subjects press at a uniform frequency throughout the trial ($8Hz$ for high, $4Hz$ for low, influence conditions), taking into account the specific guidability of the vehicle and the distance separating it from the goal. We then simply count the number of times (and divide by the number of simulated episodes, i.e. 1000) that the vehicle made it to the goal in time. In our model-based analyses of SGTs, however, we wanted to take into account possible, subtle variations in the way subjects might have perceived their pressing frequency to be higher or lower than that allowed. This "subjective prospective frequency" is then a free parameter, which we allow to vary across subjects. While models using pure probability did not account for the data most competently (and where thus discarded for DGTs analyses), for completeness (and replicability), we report our procedure to find a model-based closed form for the probability, which a sampling algorithm such as that implemented in stan can use to compute probabilities on the fly.

Thus, in SGTs, models using probability of success as a predictor generate decisions based on the estimated probability of achieving the goal. Let us call this probability $\omega(\gamma(v); d; f)$, where $\gamma(v) \in \{0.28, 0.36, 0.43\}$ is vehicle $v$'s guidability, that is, the probability that it will move in the intended direction vs. uniformly at random (we assume this value to be fully known by the time SGTs are faced); $d$ is the distance between the vehicle and its goal, and $f$ is the prospective frequency individual to each subject and influence condition.

In order to be able to make inferences about $f$ in our hierarchical Bayesian model, we built a closed form expression or approximation for the way that pressing frequency determines these probabilities (assuming optimal choices). We obtained this using the following procedure:

1. We simulated synthetic trials in which an artificial subject made optimal pressing choices at each frequency in the range $[1 : 0.1 : 10]$ (1 to 10 in steps of $0.1Hz$ ), with vehicle-goal distances in the range $[4 : 1 : 36]$ Manhattan steps, and vehicle controllabilities in the range $[0.1 : 0.025 : 0.5]$.

2. We computed the empirical probabilities for each combination of these parameters to reach the goal by running 5000 simulated trials from each.

3. Now that we have the empirical correspondences between distance, guidability, frequency of pressing, and probability of success, we can build a simple model to predict this latter as precisely as possible. We then (over)fitted a third degree multivariable polynomial (the outcome of which then goes through a logit function), to the empirical outcome probabilities; this included all three variables of vehicular controllability, distance and frequency as covariates. Thus, if for brevity we call $\gamma := \gamma(v)$ , and $f := f_{s;c}$,we have $K(\gamma; d; f)$ as our polynomial:

$$K(\gamma; d; f) = a_0 + a_1\gamma + a_2 d + a_3 f + a_4\gamma d + a_5\gamma f + a_6 df \cdots + a_{16}\gamma^3 + a_{17}d^3 + a_{18}f^3 \quad (12)$$

   The polynomial includes 19 terms in total. The probability is then computed as: $\omega(\gamma; d; f) = \sigma(K(\gamma; d; f))$ where $\sigma$ is the logistic sigmoid function.
